# GSK-3β regulates the synaptic expression of NMDA receptors via phosphorylation of phosphatidylinositol 4 kinase type IIα

**DOI:** 10.1101/841676

**Authors:** Mascia Amici, Yeseul Lee, Robert J.P. Pope, Graham L. Collingridge

**Affiliations:** Glutamate Receptor Group, School of Physiology, Pharmacology and Neuroscience, University of Bristol, Bristol, UK; Centre for Research in Neurodegenerative Disease, Department of Physiology, The University of Toronto, Toronto, Canada; Lunenfeld-Tanenbaum Research Institute, Mount Sinai Hospital, Toronto, Canada

**Keywords:** GSK-3β, PI4KIIα, rat hippocampus, NMDA receptors, long term depression

## Abstract

Understanding the normal functions of GSK-3β in the central nervous system is of major interest because deregulation of this kinase is strongly implicated in a variety of serious brain conditions, such as Alzheimer disease, bipolar disorder and schizophrenia. GSK-3β plays a role in the induction of NMDA receptor-dependent long-term depression (LTD) and several substrates for GSK-3β have been identified in this form of synaptic plasticity, including KLC-2, PSD-95 and tau. Stabilization of NMDA receptors at synapses has also been shown to involve GSK-3β, but the substrates involved are currently unknown. Recent work has identified phosphatidylinositol 4 kinase type IIα (PI4KIIα) as a neuronal GSK-3β substrate that can potentially regulate the surface expression of AMPA receptors. In the present study, we investigated the synaptic role of PI4KIIα in organotypic rat hippocampal slices. We found that knockdown of PI4KIIα had no effect on synaptic AMPA receptors but substantially inhibited synaptic NMDA receptors. Furthermore, the ability of the selective GSK-3 inhibitor, CT99021, to inhibit synaptic NMDA receptors was occluded in shRNA-PI4KIIα transfected neurons. The effects of knocking down PI4KIIα knockdown were fully rescued by a shRNA-resistant wild type construct but could not be rescued by a mutant construct that was unable to be phosphorylated by GSK-3β. The data suggest that GSK-3β phosphorylates PI4KIIα to stabilize the expression of synaptic NMDA receptors.

## INTRODUCTION

NMDA receptors are important for both synaptic transmission (Herron *et al.*, 1986) and synaptic plasticity in the hippocampus (Collingridge *et al.*, 1983). Hippocampal NMDA receptors are critically also involved in learning and memory (Morris *et al.*, 1986) and their dysregulation has been implicated in numerous brain disorders, including neurodegenerative conditions, such as Alzheimer’s disease (Snyder *et al.*, 2005), psychiatric conditions, such as schizophrenia (Mohn *et al.*, 1999; Hahn *et al.*, 2006;) and neurodevelopment conditions, such as autism (Hu *et al.*, 2016). Therefore it is important to understand how NMDA receptors are regulated at synapses.

Glycogen synthase kinase 3 (GSK-3) is a monomeric, highly conserved, multifunctional Ser/Thr kinase that was discovered for its role in glycogen metabolism (Embi *et al.*, 1980). The two paralogous proteins GSK-3α and GSK-3β are ubiquitously expressed and many studies showed that GSK-3β, in particular, is involved in a plethora of neuronal processes and disorders (Kaidanovich-Beilin & Woodgett, 2011; Bradley *et al.*, 2012). One of the key features of GSK-3β is that it is constitutively active (Hur & Zhou, 2010) and most upstream regulators act negatively to reduce GSK3 enzymatic activity via an increase of phosphorylation at its Ser9 residue. Previously we found that the activation of GSK-3β is involved in NMDA receptor-dependent long-term depression (LTD) (Peineau *et al.*, 2007), a form of synaptic plasticity that involves the endocytosis of AMPA receptors and is involved in developmental plasticity and learning and memory (Collingridge *et al.*, 2010). Subsequent work has identified some of the GSK-3 substrates involved in linking NMDA receptor activation to AMPA receptor endocytosis during LTD; these include kinesis light chain 2 (KLC2) (Du *et al.*, 2010) PSD-95 (Nelson *et al.*, 2013) and tau (Lovestone *et al.*, 1994; Takashima *et al.*, 1998; Kimura *et al.*, 2014). The regulation of AMPA receptor endocytosis may involve the guanyl nucleotide dissociation inhibitor (GDI): Rab5 complex, whereby activation of GSK-3β releases Rab5 from GDI such that Rab5 recruits AMPA receptors into early endosomes (Wei *et al.*, 2010). In addition to regulating downstream effectors of the LTD process, GSK-3β has also been shown to regulate NMDA receptor function (Chen *et al.*, 2007). This process also involves regulation of the GDI:Rab5 complex, as well as dynamin and disruption of the binding of NMDA receptors to PSD-95.

Recently phosphatidylinositol 4 kinase type II α (PI4KIIα) was identified as a neuronal GSK-3β effector with a potential role in the regulation of the surface expression of AMPA receptors (Robinson *et al.*, 2014).

In the present study, we investigated whether PI4KIIα regulates the expression of synaptic AMPA receptors and / or synaptic NMDA receptors. We used shRNA to knockdown PI4KIIα in hippocampal organotypic slices and studied synaptic transmission and LTD at the Schaffer collateral-commissural pathway. Knockdown of PI4KIIα had no effect of AMPA receptor-mediated synaptic transmission but reduced NMDA receptor mediated synaptic transmission. Pharmacological inhibition of GSK-3 similarly reduced NMDA receptor-mediated synaptic transmission; an effect that was precluded by prior knockdown of PI4KIIα. The effects of PI4KIIα knockdown on NMDA receptor-mediated synaptic transmission were fully rescued by expression of wildtype PI4KIIα but not by expression of a mutant form of PI4KIIα that could not be phosphorylated by GSK-3β. These results suggest that GSK-3β phosphorylates PI4KIIα to stabilize NMDA receptors at synapses.

## Materials and methods

### Organotypic hippocampal slices

Transverse hippocampal slices from p8 Wistar rats were prepared as described previously (Rocca *et al.*, 2013). They were cut in ice-cold aCSF (in mM: 238 sucrose, 2.5 KCl, 26 NaHCO_3_, 1 NaH_2_PO_4_, 1 CaCl_2_, 9 MgSO_4_, 10 glucose) saturated with 95% O_2_/ 5% CO_2_. Slices were placed on Millicell culture plate inserts (Merck Millipore) and maintained at 35°C, 5% CO_2_in MEM based culture media containing 20% horse serum and (in mM): 30 HEPES, 16.25 glucose, 5 NaHCO_3_, 1 CaCl_2_, 2 MgSO_4_, 0.68 ascorbic acid and 1μg/ml insulin, pH 7.28, 320 mOsm.

### Biolistic transfection

Organotypic slices were biolistically transfected at 3-5 days *in vitro*(div) using a Helios GeneGun (Bio-Rad), bullets were prepared using PI4KIIα shRNA alone or in combination with either wild type or S9/51A shRNA resistant PI4KIIα. GFP was used as a transfection marker.

### Electrophysiology

Whole cell voltage-clamp recordings were made from CA1 pyramidal cells at 6-11 div, blind with respect to the transfected plasmid. Patch pipettes contained intracellular solution (in mM): 8 NaCl, 130 Cs-methanesulfonate, 10 HEPES, 0.5 EGTA, 4 MgATP, 0.3 Na_3_GTP, 5 QX-314, pH 7.25, 290 mOsm. Picrotoxin (50 μM) and 2-chloroadenosine (1 μM) were routinely included in the bath solution, which comprised (in mM): 124 NaCl, 3 KCl, 26 NaHCO_3_, 1.4 NaH_2_PO_4_, 4 CaCl_2_, 4 MgSO_4_, 10 glucose; saturated with 95% O_2_/ 5% CO_2_). In order to isolate NMDA receptor-mediated EPSCs (NMDAR-EPSCs), 3 μM NBQX was added and neurons depolarized to −40 mV; in some cases, D-AP5 (50 μM) was added at the end of the experiment to confirm synaptic responses were NMDAR mediated. Two stimulating electrodes (test and control input) were placed in the Schaffer collateral-commissural pathway and stimulated at 0.05 Hz to record AMPAR-EPSCs and at 0.03 Hz for NMDAR-EPSCs. Long-term depression (LTD) was induced using a pairing protocol: 1Hz for 6 min, Vh = −40mV. Data were acquired and analysed with WinLTP (Anderson & Collingridge, 2007). Statistical analysis was performed using paired or unpaired Student’s *t* test or one-way ANOVA as appropriate, and significance was set at p < 0.005.

## RESULTS

### PI4KIIα knock down reduced expression of synaptic NMDA receptors

To determine whether PI4KIIα regulates the synaptic expression of either AMPA or NMDA receptors we utilized three previously validated plasmids (Robinson *et al.*, 2014). These were an shRNA probe to reduce the expression of PI4KIIα and two shRNA resistant constructs for rescue experiments; a wild-type (WT) and a mutant PI4KIIα (S9/51A) that cannot be phosphorylated by GSK-3β. CA1 pyramidal neurons were biolistically transfected with these constructs (plus GFP) and dual whole-cell patch-clamp electrophysiological recordings obtained in response to stimulation of the Schaffer collateral-commissural pathway.

Knockdown of PI4KIIα had no effect on AMPA receptor-mediated excitatory post synaptic currents (EPSC-A) as ascertained by comparing the amplitude of the evoked synaptic response in a transfected neuron and a nearby non-transfected neuron, at a holding potential of −70 mV (Fig 1A). In contrast, in the same neurons recorded at +40 mV, knockdown of PI4KIIα led to a significant reduction in the EPSC recorded at a latency of 60 ms, at a time when the response is dominated by an NMDA receptor-mediated component (EPSC-N) (Fig 1B). To obtain a more accurate estimate of the reduction in EPSC-N, we pharmacologically isolated the pure NMDA receptor-mediated synaptic current by blocking the AMPA receptor-mediated component with NBQX (3μM). We made recordings at −40 mV and observed a substantial reduction in the amplitude of EPSC-N in transfected compared to non-transfected pairs of neurons (Fig 1C). The effect of the shRNA construct could be ascribed to knockdown of PI4KIIα since there was full rescue in neurons in which an shRNA-resistant wild type PI4KIIα was co-transfected with the shRNA-PI4KIIα (Fig 1D). In contrast, a mutated shRNA resistant PI4KIIα that could not be phosphorylated by GSK-3 (S9/51A) was not able to rescue the reduction in EPSC-N (Fig 1E). Quantification of the effects of these transfections on EPSC-N is shown in Figure 1F.

**Figure 1.**
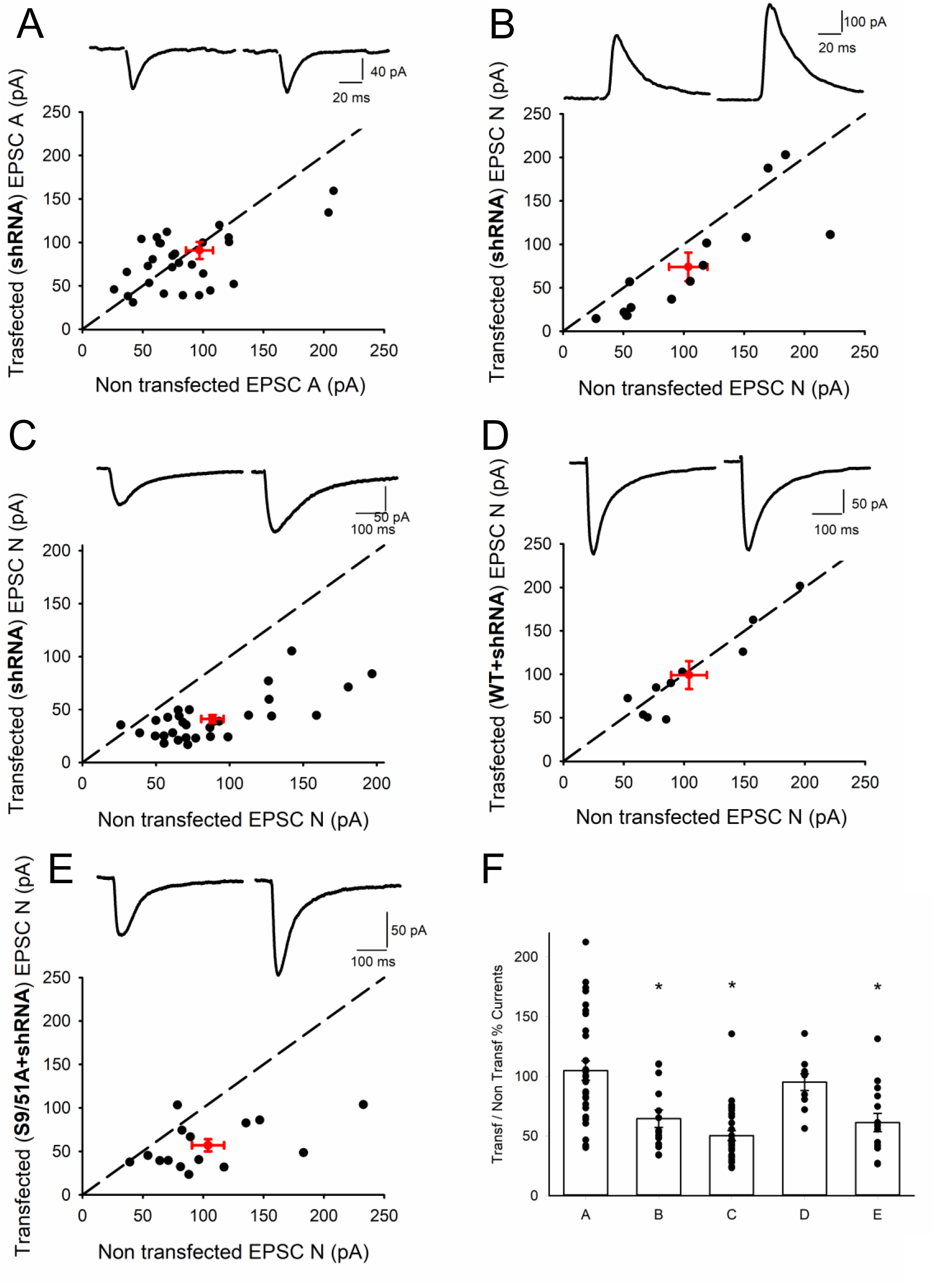
PI4KIIα knock down reduces NMDA receptor-mediated synaptic transmission. In graphs A-E EPSCs were recorded in transfected cells and nearby non-transfected cells, amplitudes were plotted for each pair (black circles), red circle represents mean ± sem. Insets show representative traces (left: transfected; right: non-transfected cell). **A)** EPSC-As were recorded in transfected cells (shRNA PI4KIIα) and nearby non-transfected cells. Transfected cells −91±10 pA, Non-transfected −97±11 pA (n=31). **B)** EPSC-Ns were measured 60 ms post-stimulation (at +40 mV). Transfected cells 74±16 pA, Non transfected 104±16 pA (n=14). **C)** In a different set of cells peak amplitudes in the presence of 3 μM NBQX (at −40 mV) were plotted for each pair. Transfected cells −41±4pA, Non transfected −88±8 pA (n=30). **D)** Pharmacologically isolated EPSC-Ns were measured in cells transfected with shRNA and WT PI4KIIα (−99±16pA) and nearby non-transfected cells (−104±15 pA n=10). **E)** Pharmacologically isolated EPSC-Ns were measured in cells transfected with shRNA and S9/51A PI4KIIα (−57±7pA) and nearby non transfected cells (−104±13 pA n=15). **F)** Summary showing the mean and data points of the ratio of the current amplitude in transfected over paired non-transfected cell for each condition, * indicates the presence of a statistically significant difference between transfected and non-transfected cells within a group (labelled after the panels in this figure) following a paired Student’s *t*-test.

### PI4KIIα is required for a GSK-3 inhibitor to inhibit EPSC-N

Previous studies have shown that inhibition of GSK-3 results in the internalization of NMDARs (Chen *et al*, 2007). Our observation that knockdown of PI4KIIα had a similar effect suggest that PI4KIIα may be a downstream effector of GSK-3 in the regulation of synaptic NMDA receptors. To test this directly we inhibited GSK-3 using CT99021, its most selective inhibitor (Bain *et al.*, 2007), in control and transfected neurons. CT99021 (1μM) inhibited EPSC-N in control neurons but had no effect in neurons transfected by the shRNA-PI4KIIα construct (Fig 2A). The ability of CT99021 to inhibit EPSC-N was rescued in neurons transfected with the wild type PI4KIIα construct but not with the S9/51A mutant construct (Fig 2B). These data, which are quantified in Figure 2C, suggest that GSK-3β phosphorylates PI4KIIα to stabilize synaptic NMDA receptors.

**Figure 2.**
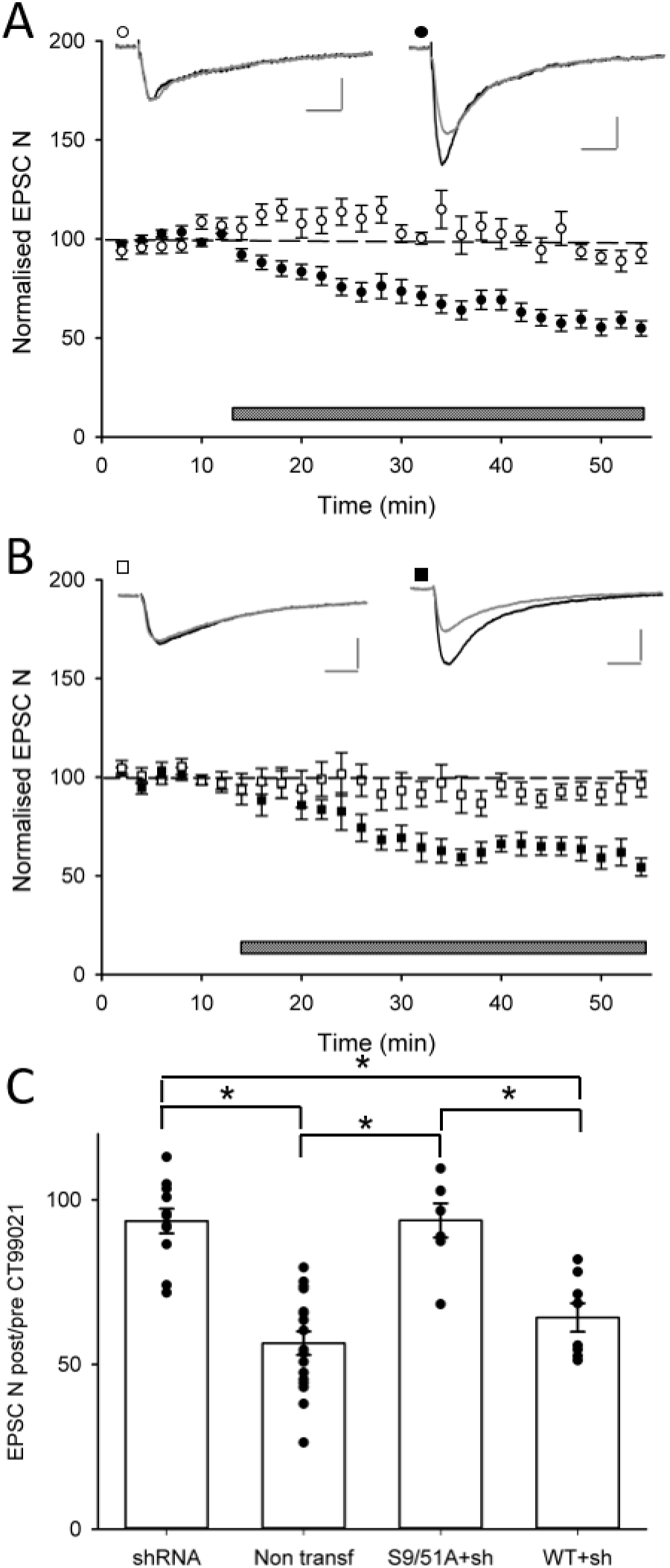
GSK-3β stabilizes synaptic NMDA receptors via phosphorylation of PI4KIIα. **A** Application of a GSK-3 inhibitor (CT99021, 1μM) reduced EPSC-N in non-transfected cells (black circles n=17), but not in cells transfected with shRNA to knockdown PI4KIIα (white circles n=11). **B** The effect of CT99021 was rescued when wild type PI4KIIα shRNA resistant was co-expressed with the shRNA (black squares n = 8) but not when the mutated S9/51A shRNA resistant PI4KIIα was co-expressed with the shRNA (white squares n = 7). **C** Summary histograms of the effects of CT999021 (quantified 30 min after start of application) for the various conditions. * indicates the presence of a statistically significant difference between groups. Insets show representative traces before (black) and 30’ after (grey) application of CT99021. Calibration bar: 50pA/50 ms.

### PI4KIIα is not required for NMDAR dependent LTD

Our observations show that inhibition of PI4KIIα results in approximately a 50% reduction in the synaptic expression of NMDA receptors. Since NMDA receptors are required for the induction of LTD at these synapses we wondered whether inhibition of PI4KIIα would impair this process or whether the residual synaptic NMDA receptors could support LTD. To test this possibility, we performed two pathways experiments, where we used a pairing protocol to induce LTD in the test pathway while leaving unperturbed the control pathway. We used the same induction protocol when recording EPSC-A and EPSC-N (Rocca *et al.*, 2013). Neither LTD of EPSC-A or EPSC-N were affected by knocking down PI4KIIα (Figure 3). Therefore, the residual synaptic NMDA receptors are sufficient to enable synaptic plasticity.

**Figure 3.**
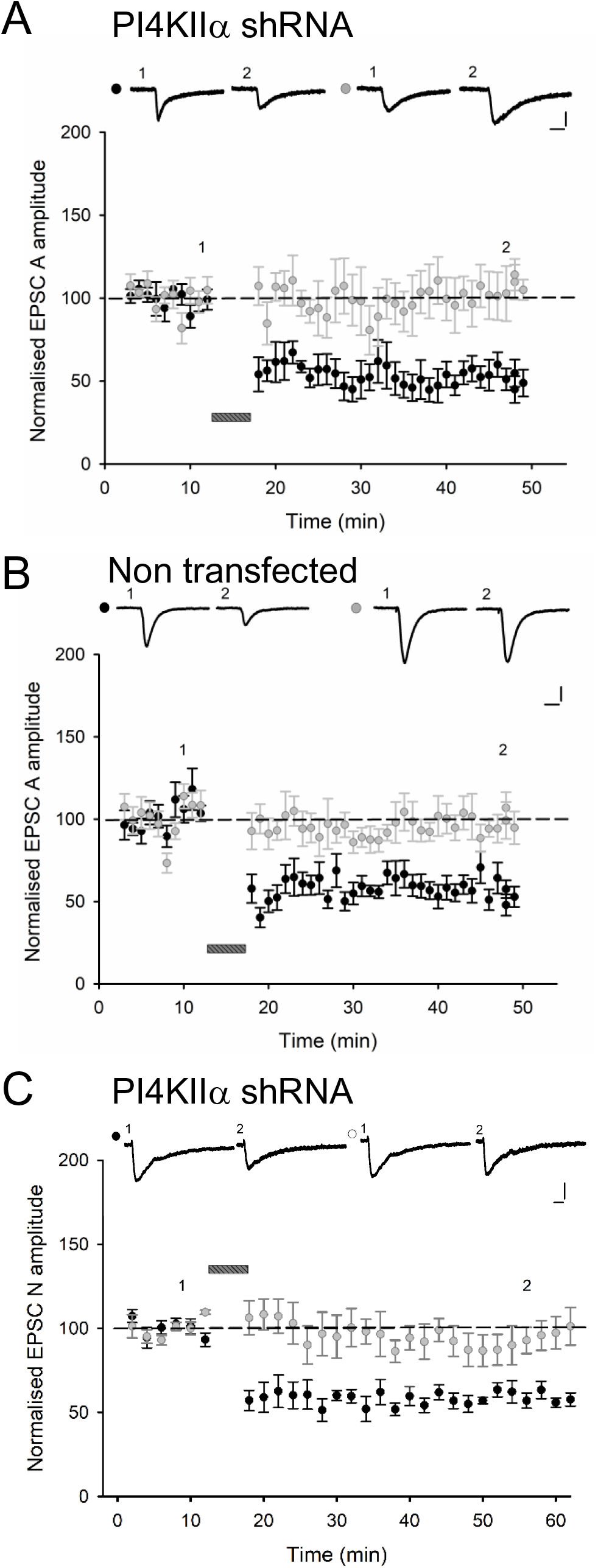
PI4KIIα is not required for NMDAR-dependent LTD. **A,B** Similar levels of input-specific LTD was recorded in cells transfected with shRNA-PI4KIIα (**A**) and non-transfected (**B**) cells. The bar indicates delivery of a pairing protocol (1 Hz for 6 min, Vh = −40 mV). Representative traces for test (black circle) and control (grey circle) pathway at time points indicated; calibration bar 50pA/20ms. Depression, quantified 30 min after the pairing protocol, was similar in both sets of cells: 54 ± 4% (n = 8) and 52 ± 7 % (n = 7) of baseline, respectively. **C** Similar levels of input specific LTD of EPSC-N was recorded in cells transfected with shRNA PI4KIIα and control cells. Depression, quantified 30 min after the pairing protocol, (61±4% of baseline n=4) is comparable to that obtained in non transfected cells (not shown), calibration bar: 30pA/50ms.

## Discussion

In the present study we have provided evidence that PI4KIIα is required to sustain a full synaptic complement of NMDA receptors. In neurons in which PI4KIIα had been knocked-down we found that inhibition of GSK-3 no longer affected NMDA receptor-mediated synaptic transmission. Furthermore, we found that this effect was fully rescued by wildtype PI4KIIα but not by a mutant form of PI4KIIα that could not be phosphorylated by GSK-3β. The most straightforward explanation for these results is that GSK-3β phosphorylates PI4KIIα to maintain a full synaptic complement of NMDA receptors.

### PI4KIIα and the regulation of AMPA receptors

We were surprised that knockdown of PI4KIIα had no effect on AMPA receptor-mediated synaptic transmission given previous work, using the same constructs, found a modest increase in cell surface AMPARs in cultured hippocampal neurons (Robinson *et al.*, 2014). There are a number of possible explanations for this difference. First, we studied only synaptic AMPARs whereas the previous study measured the total complement of cell surface GluA1, which labels both synaptic and extra synaptic AMPA receptors (Richmond *et al.*, 1996). Indeed, GluA1 homomers are preferentially located at extrasynaptic sites and are only driven into these synapses transiently during certain forms of synaptic plasticity (Park *et al.*, 2016). We wondered, however, whether an excess of surface GluA1 might result in the appearance of GluA2-lacking, calcium permeable AMPARs at synapses transfected by the shRNA construct. However, there was no difference in the rectification of EPSC-A in transfected and non-transfected neurons (unpublished observations). A second possibility is that we had not knocked down PI4KIIα sufficiently to affect AMPA receptor trafficking. Either way, our results clearly show that the dominant effect of knocking down PI4KIIα is to regulate the synaptic complement of NMDA receptors.

### PI4KIIα and the regulation of NMDA receptors

Previous work had shown that inhibition of GSK-3 leads to a reduction in the cell surface of NMDA receptors, including NMDA receptor-mediated synaptic currents (Chen *et al*, 2007). Our work confirms, using a more specific GSK-3 inhibitor, CT99021, this effect. The finding that knockdown of PI4KIIα occludes the effect of CT99021 suggests that GSK-3 is operating via this lipid kinase to mediate this action. Furthermore, the finding that the effect of the knockdown by PI4KIIα on ECPC-N is fully rescued by wild type PI4KIIα but not by a mutant, in which the priming sites (Ser9 and Ser51) for GSK-3β were mutated to alanine, strongly suggest that GSK-3β phosphorylates PI4KIIα to regulate synaptic NMDA receptor function. In terms of the underlying mechanism, there is evidence that constitutive GSK-3β activity maintains NMDA receptors on the neuronal plasma membrane by inhibiting their internalization. This clathrin and dynamin dependent internalization process involves a Rab5-directed transport of endocytic vesicles to early endosomes (Chen *et al*, 2007). It seems likely therefore that PI4KIIα is part of the machinery that links GSK-3 activity to this intracellular pathway for NMDA receptor removal and degradation. PI4KIIα is located on the plasma membrane, on endosomes and the trans-Golgi network. PI4KIIα is also cargo for adaptor protein complex 3 (AP3) and is capable of regulating the sorting from endosomes to lysosomes (Salazar *et al.*, 2005; Craige *et al.*, 2008), Minogue, 2018). Indeed, it has been proposed that a major role of PI4KIIα is to bind to AP-3 on endosomes and catalyse the local formation of PtdIns(4)P, which then recruits additional PI4KIIα and other proteins involved in the formation and targeting of AP-3 vesicles (Craige *et al.*, 2008). One possibility therefore is that PI4KIIα drives the AP3-mediated trafficking of proteins, such as dysbindin or the chloride channel CLC-3 (Farmer *et al.*, 2013), that limit the functional expression of NMDA receptors (Tang *et al.*, 2009, Jeans *et al.*, 2011). Alternatively, it may regulate components of the endocytotic pathway that removes NMDA receptor subunits from the plasma membrane. In this context, dysbindin is thought to be involved in trafficking in the lysosomal pathway and its knockout leads to the accumulation of GluN2A subunits and potentiation of NMDA receptor-mediated synaptic transmission (Tang et al, 2009). However, there is also good evidence that PI4KIIα has a role in driving traffic toward the plasma membrane (Minogue, 2018). This may involve the hydrolysation of PtdIns3P in early/sorting endosomes to PtdIns substrates such that PI4KIIα can produce PtdIns4P and the recruitment of the exocyst complex. How precisely PI4KIIα acts to maintain the full complement of NMDA receptors remains to be determined.

### PI4KIIα and the regulation of LTD

An interesting observation was that despite a substantial reduction in the synaptic expression of NMDA receptors, by ~50%, NMDA receptor-mediated LTD was unaffected by PI4KIIα knockdown. Firstly, this shows that PI4KIIα is unlikely to be part of the LTD machinery that links GSK-3β to AMPA receptor internalization and hence distinguishes it from other GSK-3β substrates that have been directly implicated in this process, such as KLC2 (Du *et al.*, 2010), PSD-95 (Nelson *et al.*, 2013) and tau (Kimura *et al.*, 2014). Secondly, it also shows that inhibition of NMDA receptors by GSK-3 antagonists is unlikely to account for the ability of these antagonists to block the induction of LTD; rather they block the ability of GSK-3 to regulate the AMPA receptor endocytic machinery directly (Peineau *et al.*, 2007). Thirdly, it suggests the existence of two pools of synaptic NMDA receptors, one regulated by GSK-3β and PI4KIIα and another that is not; the latter being able to support the induction of NMDA receptor dependent LTD. Of potential relevance is the finding that dysbindin regulates the internalization of a GluN2A-containing pool of NMDA receptors that is involved in LTP but not LTD (Tang et al, 2009). Speculatively, PI4KIIα might selectively stabilize this pool of GluN2A-containing NMDA receptors. A consequence of this action would be to preserve the ability of synapses that have undergone LTD, and hence have a reduced number of synaptic NMDA receptors, to still be capable of exhibiting NMDA receptor-dependent LTP. This mechanism would accordingly preserve the bidirectional modifiability of synaptic strength, in the face of a reduction in synaptic NMDA receptors.

It is interesting to note that synaptic AMPA receptors are also composed on a constitutive recycling pool that is stabilized by an interaction between NSF and the GluA2 subunit and a stable pool that is not regulated in this manner (Nishimune *et al.*, 1998). Further work is required to understand the significance and underlying mechanisms that regulate these two synaptic pools of NMDA receptors.

## Acknowledgements

We would like to thank Dr Adam Cole and his group (Garvan Institute of Medical Research, Sydney, Australia) for providing us with the plasmids used in this study.

## Funding

This work was supported by the MRC, ERC (UK) and CIHR (Canada).

## Conflict of Interest Statement

The authors declare no conflict of interests.

## Author Contributions

MA and GLC conceived the project; MA designed and carried out all the experiments and performed the analysis. YL, RJPP contributed to the design of experiments. MA and GLC wrote the paper.

## Data Accessibility Statement

Data are available upon request from the corresponding author.

## Abbreviations

AMPA: α-amino-3-hydroxy-5-methyl-4-isoxazole propionic acid
EPSC: excitatory post synaptic current
GSK-3β: glycogen synthase kinase 3 beta
LTD: long term depression
NMDA: *N*-methyl-*D*-aspartate
PI4KIIα: phosphatidylinositol 4 kinase type II α

